# Ribosome-targeting antibiotics impair T cell effector function and ameliorate autoimmunity by blocking mitochondrial protein synthesis

**DOI:** 10.1101/832956

**Authors:** L Almeida, A Dhillon-LaBrooy, CN Castro, N Ayele, J Bartel, GM Carriche, M Guderian, S Lippens, S Dennerlein, C Hesse, BN Lambrecht, L Schauser, BR Blazar, M Kalesse, R Müller, LF Moita, T Sparwasser

## Abstract

While antibiotics are intended to specifically target bacteria, most are known to affect host cell physiology. In addition, some antibiotic classes are reported as immunosuppressive, for reasons that remain unclear. Here we show that linezolid, a ribosomal-targeting antibiotic (RAbo), effectively blocked the course of a T cell-mediated autoimmune disease. Linezolid and other RAbos were strong inhibitors of Th17 effector function *in vitro*, showing that this effect was independent of their antibiotic activity. Perturbing mitochondrial translation in differentiating T cells, either with RAbos or through the inhibition of mitochondrial elongation factor G1 (mEF-G1) progressively compromises the integrity of the electron transport chain (ETC). Ultimately, this leads to loss of mitochondrial metabolism and cytokine production in differentiating Th cells. In accordance, mice lacking *Gfm1* in T cells are protected from EAE, demonstrating that this pathway plays a key role in maintaining T cell function and pathogenicity.

## Introduction

Bacteria require intact cell walls and functional ribosomes to retain their structural and metabolic integrity. As such, inhibition of these processes by antibiotics either kills bacteria or prevents their growth and division (1–3). As eukaryotic cells do not possess cell walls and their ribosomes are structurally different from bacterial ribosomes, antibiotic drugs can be used in the clinic to eradicate bacterial infections, with limited toxicity to the host. Nevertheless, some antibiotics have reported side effects, such as candidiasis, anemia, bone marrow suppression, lactic acidosis and immunosuppression (4–11). Since antibiotics are often administered to patients who have ongoing bacterial infections, it is important to understand the mechanisms governing antibiotic-mediated immunosuppression. Without a proper host immune response, both clearance of infections and control of opportunistic pathogens are impaired. Strikingly, both the cellular and molecular mechanisms governing immunosuppression by antibiotics are poorly understood. Here, we focus on linezolid, a ribosomal-targeting antibiotic (RAbo) from the Oxazolidinone family which targets Gram-positive bacteria (12) and has been approved (and is reserved) for treatment of several multi-resistant bacterial infections (13). Patients requiring linezolid treatment often present in a critical state and are at increased risk of fungal infections, especially after prolonged administration (14). Moreover, linezolid is associated with higher mortality by Gram-negative pathogens when compared with vancomycin. Both antibiotics share a similar spectrum of action, and therefore this discrepancy is unusual (15).

Linezolid, unlike vancomycin, pertains to the class of RAbos, which have been shown to inhibit mitochondrial translation in mammalian systems and cell lines (16–18). It achieves this by occupying the peptidyl transferase center at the 50S subunit of the mitochondrial ribosome, interfering with the binding of aminoacyl-tRNAs (19). The correct assembly of electron transport chain (ETC) complexes relies on the coordinated translation of proteins which are simultaneously derived by cytosolic and mitochondrial ribosomes (20, 21). By blocking mitoribosomes, RAbos can lead to an imbalance of nuclear and mitochondrial-encoded ETC subunits (mitonuclear imbalance) (22). The concept of mitonuclear imbalance and its relevance to organismal physiology has been previously explored in different fields of medical research. Specifically, inhibition of mitochondrial translation, which leads to a decrease in mitochondrial-encoded subunits, affects adipokine secretion in adipocytes (23), improves worm lifespan in *C. elegans* (22), and selectively kills leukaemia cells (24). The consequence of blocking mitochondrial protein synthesis in immune cells however, has not been elucidated.

Here, we show that linezolid exhibits strong immunosuppressant properties, as illustrated by its capacity to disrupt the development of self-reactive Th cells in a mouse model of multiple sclerosis. Linezolid and other RAbos were additionally able to directly inhibit Th17 function *in vitro*, showing that immunosuppression occurs independently of their antibiotic activity. Mechanistically, we demonstrate that inhibitors of mitochondrial translation lead to a mitonuclear imbalance in differentiating Th17 cells, which culminates in loss of mitochondrial activity and cytokine production. In conclusion, administration of RAbos should be carried out with extra caution to prevent undesired immunosuppression. On the other hand repurposing of RAbos and/or development of novel drugs with a similar mechanism of action, could potentially be used to control T-cell mediated autoimmune disorders.

### Linezolid inhibits onset of a T cell mediated autoimmune disease

To determine if Th17 cell responses *in vivo* could be altered by RAbos treatment, we explored a murine model of T-cell mediated autoimmunity, namely Experimental Autoimmune Encephalomyelitis (EAE). MOG-immunized mice develop severe, classical, clinical signs of EAE, presenting with infiltrating Th cells in the central nervous system (CNS), as well as decreased motility. However, administration of a human-equivalent dose of linezolid limited the development and severity of EAE symptoms (**Fig. 1a, b**). This correlated with lower frequencies and numbers of infiltrating Th cells in the CNS of linezolid-treated mice (**Fig. 1c**). Linezolid treatment led to a marked decrease in MOG-specific IL-17A- and IFN-γ-producing Th cells, without effects on the frequency of Foxp3^+^ Th cells. Oral antibiotics can lead to disturbances of microbial populations in the intestinal lumen, which in turn can influence the outcome of an immune response (25–27). To rule out that any effects mediated by linezolid were due to its antibacterial activity, vancomycin was used as an additional control, which holds a similar spectrum of activity against bacteria as linezolid. Vancomycin-treated mice developed severe EAE symptoms comparable to those of the untreated group, strongly suggesting that linezolid’s immunosuppression was due to its effect on host cells, and not bacterial cells.

**Figure 1:**
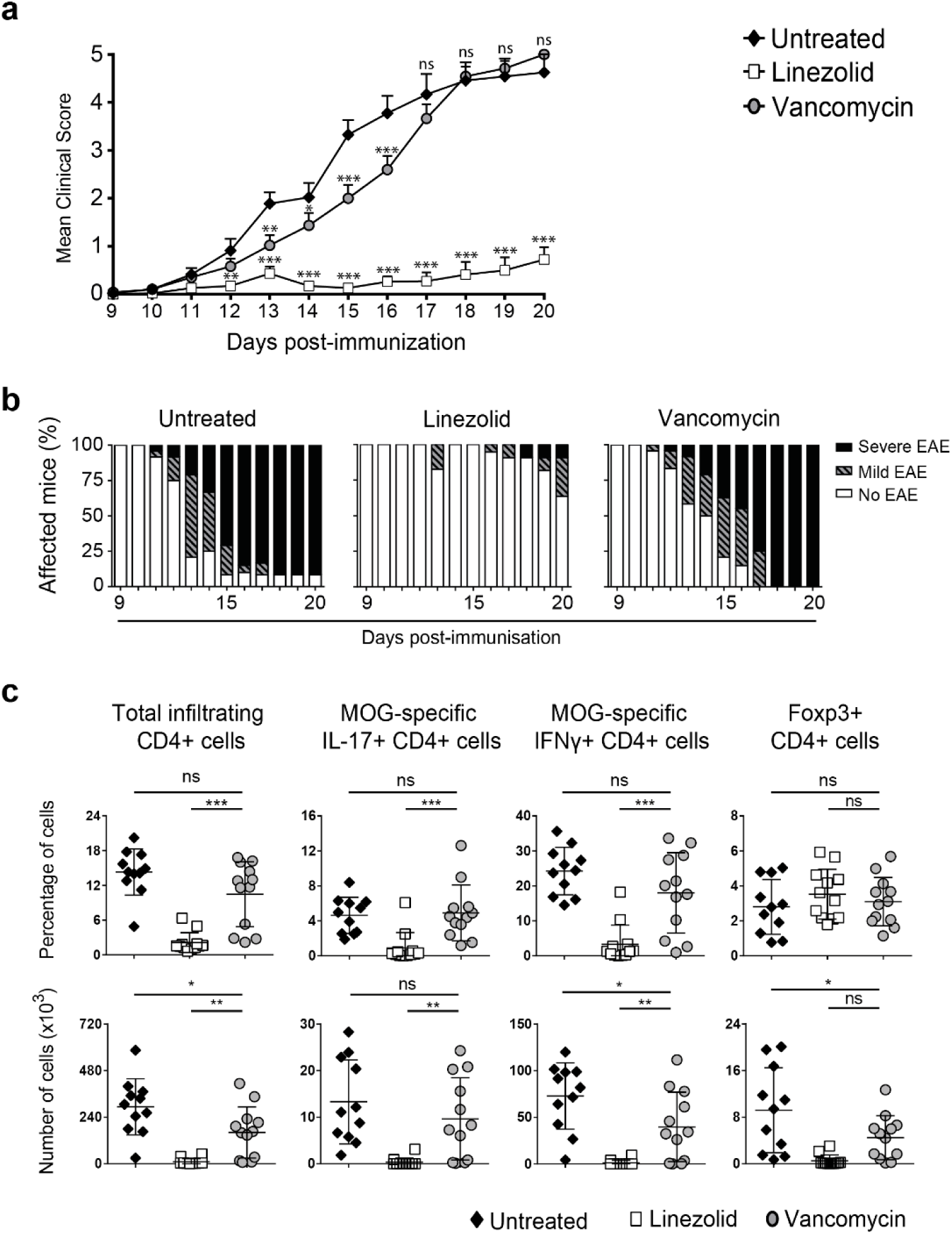
Linezolid inhibits the onset of T cell mediated autoimmune diseases EAE, in MOG-immunized mice. **(a-c)** Mice were immunized with MOG_35-55_ in CFA and pertussis toxin to induce EAE. **(a)** EAE clinical score according to clinical symptoms of mice treated daily with linezolid, vancomycin or untreated. **(b)** Distribution of disease severity: No EAE: score < 1 (□), mild EAE: 1 ≤ score < 3 (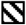), severe EAE: score ≥ 3 (▪). **(c)** Frequency (top row) and total numbers (bottom row) of CD4^+^ T cells isolated from the central nervous system (brain and spinal cord) of linezolid-, vancomycin-treated and untreated mice. Plots are obtained from the pooled data of three **(a-c)** individual experiments, with error bars showing the S.E.M of the pooled scores **(a)**. Statistical significance was determined using 2-way ANOVA with Bonferroni multiple corrections test compared to “untreated” mice.

### Mitochondrial translation inhibition by antibiotics inhibits Th17 cell function

Linezolid was able to strongly inhibit EAE induction when compared to vancomycin, which prompted us to look at differences in their mechanism of action. While vancomycin inhibits cell wall synthesis in bacterial cells, linezolid works by inhibiting prokaryotic ribosomes, blocking bacterial protein synthesis. Because mitochondria are of prokaryotic origin, there are substantial similarities between bacterial and mitochondrial mRNA translation machinery (28). Consequently, some RAbos can also inhibit mitoribosomes. This is true not only for linezolid, but also for other antibiotics pertaining to different classes such as tigecycline (a Glycylcycline) and thiamphenicol (an Amphenicol) (29–31). Hence, we postulated that RAbos inhibited Th17 function independently of their antibiotic function, through inhibition of mitochondrial translation. In accordance, when murine naïve CD4^+^ T cells were cultured under Th17 polarizing conditions in the presence of the aforementioned antibiotics, their IL-17 production was reduced by all three drugs in a dose-dependent manner **(Fig. 2a**, above). Importantly, loss of cytokine-producing cells was not caused by death of effector cells, as viability was mostly unaffected **(Fig. 2a**, below). To verify that this effect was not exclusive to murine cells, we differentiated human cord blood-derived naïve T cells under Th17-polarizing conditions in the presence of linezolid. Accordingly, human Th17 cells equally presented a lower IL-17A secretion upon linezolid treatment **(Fig. 2b**, left). Like in the murine system, viability was not affected at the indicated concentrations **(Fig. 2b**, right). We sought to determine if linezolid was able to acutely inhibit mitochondrial translation in T cells. To this end, we labeled mitochondrial translation products with [^35^S]-methionine in a murine lymphoma T cell line (EL-4), pretreated for 1h with linezolid. We observed a near complete blockage of [^35^S]-methionine incorporation into mitochondrial-encoded ETC subunits by linezolid at 100µM (**Fig. 2c**), which correlated with the concentration required to fully inhibit IL-17 production (**Fig. 2a**). The ETC is composed of five complexes, made up of several protein subunits, which must be correctly assembled in the inner mitochondrial membrane. While all complex II subunits are encoded in the nucleic DNA (nDNA) and translated in the cytosol, complexes I, III, IV and V contain subunits encoded by both nDNA and mitochondrial DNA (mtDNA). Therefore, the assembly of these complexes requires a tight coordination between the cytosolic and mitochondrial protein machinery. When mitochondrial translation is exogenously interrupted, a mitonuclear imbalance occurs, as illustrated by a higher ratio of nDNA to mtDNA-encoded subunits (16, 22). Accordingly, the presence of linezolid leads to the selective depletion of mtDNA-encoded mitochondrial-NADH-ubiquinone oxidoreductase chain 1 (ND1) and cytochrome C oxidase I (COX1) (**Fig. 2d**). By contrast, the levels of nuclear-encoded succinate dehydrogenase complex flavoprotein subunit A (SDHA) remain unaffected (**Fig. 2d**). These findings confirm that mitoribosomes – but not cytosolic ribosomes – are targets of linezolid.

**Figure 2:**
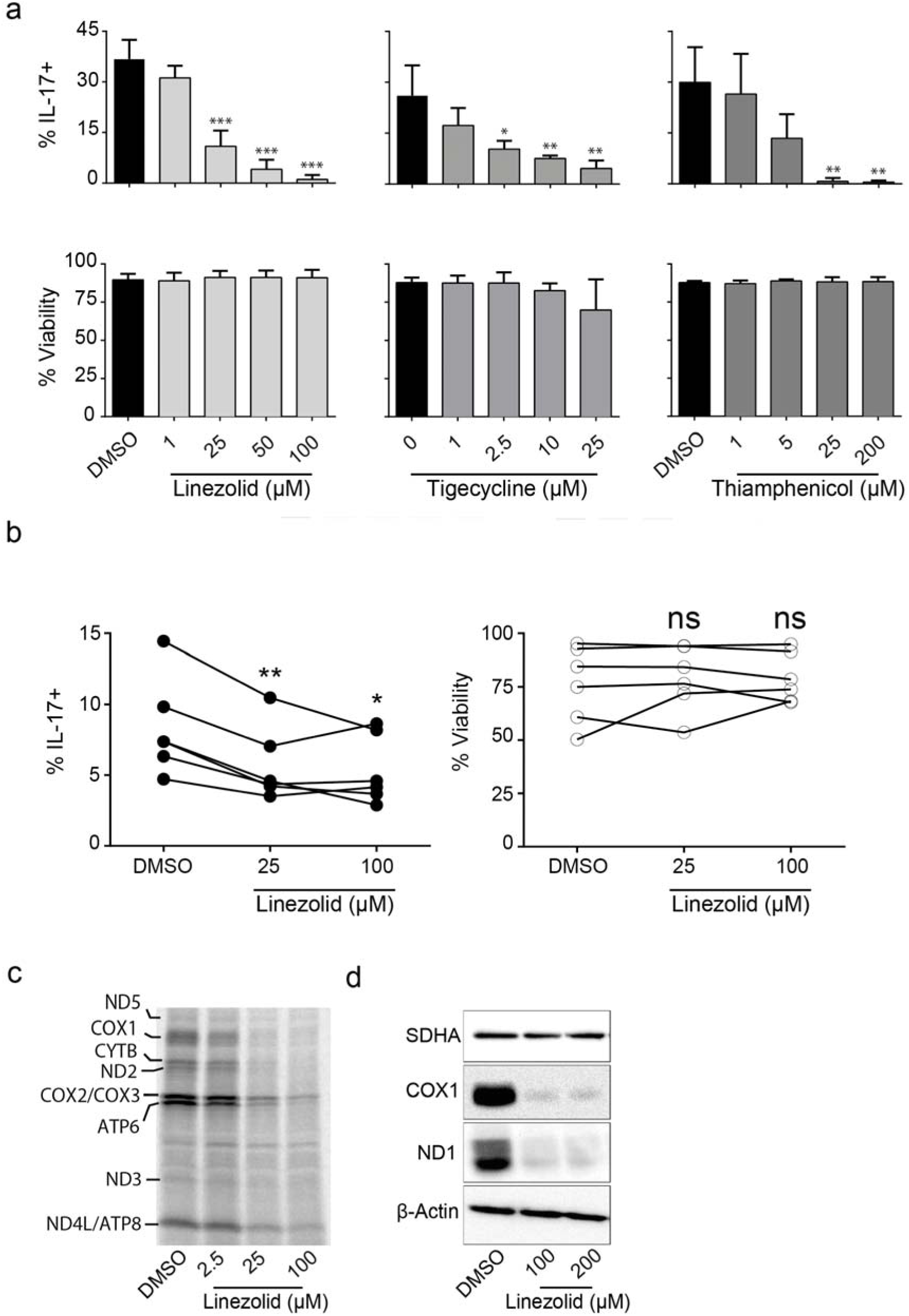
Linezolid and other RAbos disrupt Th17 effector function by targeting mitochondrial translation. **(a-d)** Naïve CD4^+^ T cells were cultured under Th17 polarizing conditions in the presence of the indicated concentrations of linezolid, tigecycline and thiamphenicol. **(a)** After 96h of culture, cells were stained for intracellular IL-17A. Bar graphs represent % of IL-17A^+^ cells amongst live CD4^+^ cells (top), and bar graphs showing % viable CD4^+^ cells (bottom). **(b)** Human naïve T cells were stained for intracellular IL-17A (right), after being differentiated for 6 days in the presence of the indicated concentrations of linezolid. Graphs contain the data for each of 6 individual donors, indicated by dots connected by the same line. The values represent % of IL-17A^+^ cells amongst live CD4^+^ cells (left) and the % of viable CD4^+^ cells (right). **(c)** Representative plot showing mitochondrial translation levels of EL-4 cells pre-incubated for 1h with linezolid or DMSO and radioactively labeled with [^35^S]-methionine. **(d)** Representative plot showing total protein levels of SDHA, COX1 and ND1, measured by western blot after 96h of culture. Plots are representative of two **(c)** or three **(d)** experiments and bar graphs are pooled means of technical replicates from three **(a**, middle, right and **c)**, or four **(a**, left) independent experiments.

### Argyrin C and linezolid exert analogous effects on differentiating Th17 cells

Mitochondrial ribosome subunits are encoded by the *MT-RNR1* and *MT-RNR2* genes, which encode the 12S and 16S ribosomal RNA (rRNA), respectively. These rRNAs differ between individuals, depending on their mitochondrial haplotype (32–34). This has the potential to modulate the susceptibility of mitoribosomes to RAbos, as shown previously for linezolid (35). To explore different ways of inhibiting mitochondrial translation, we employed the Argyrin (Arg) family of compounds (**Fig. 3a** and **S1a**). Argyrins do not bind to the mitoribosome, but instead to mitochondrial elongation factor G1 (mEF-G1) (36), an enzyme necessary for the elongation step during mitochondrial protein synthesis (37). The inhibitory capacities of the different Arg family members on Th17 cytokine production were quantified in cultured murine naïve CD4^+^ T cells under Th17 polarizing conditions (**Fig. S1a, b**). Arg A and B effectively inhibited the frequency of IL-17A-producing T cells only at the highest concentration (240nM) tested, while Arg C and D were effective at much lower doses (24nM). Notably, Arg F and G had no immunosuppressive activity at the tested concentrations despite holding high homology with the most active family members Arg C and D (**Fig. S1b**, left). Importantly, Arg F still retained the capacity to inhibit IL-17 production, albeit at higher concentrations, (>600nM) (**Fig. S1c**, left). This is consistent with a previous report, showing that Arg F inhibited COX2 expression at doses higher than 600nM (38). As we observed for RAbos, cellular viability was not affected by any Argyrin (**Fig. S1b**, right and **S1c**, right).

**Figure 3:**
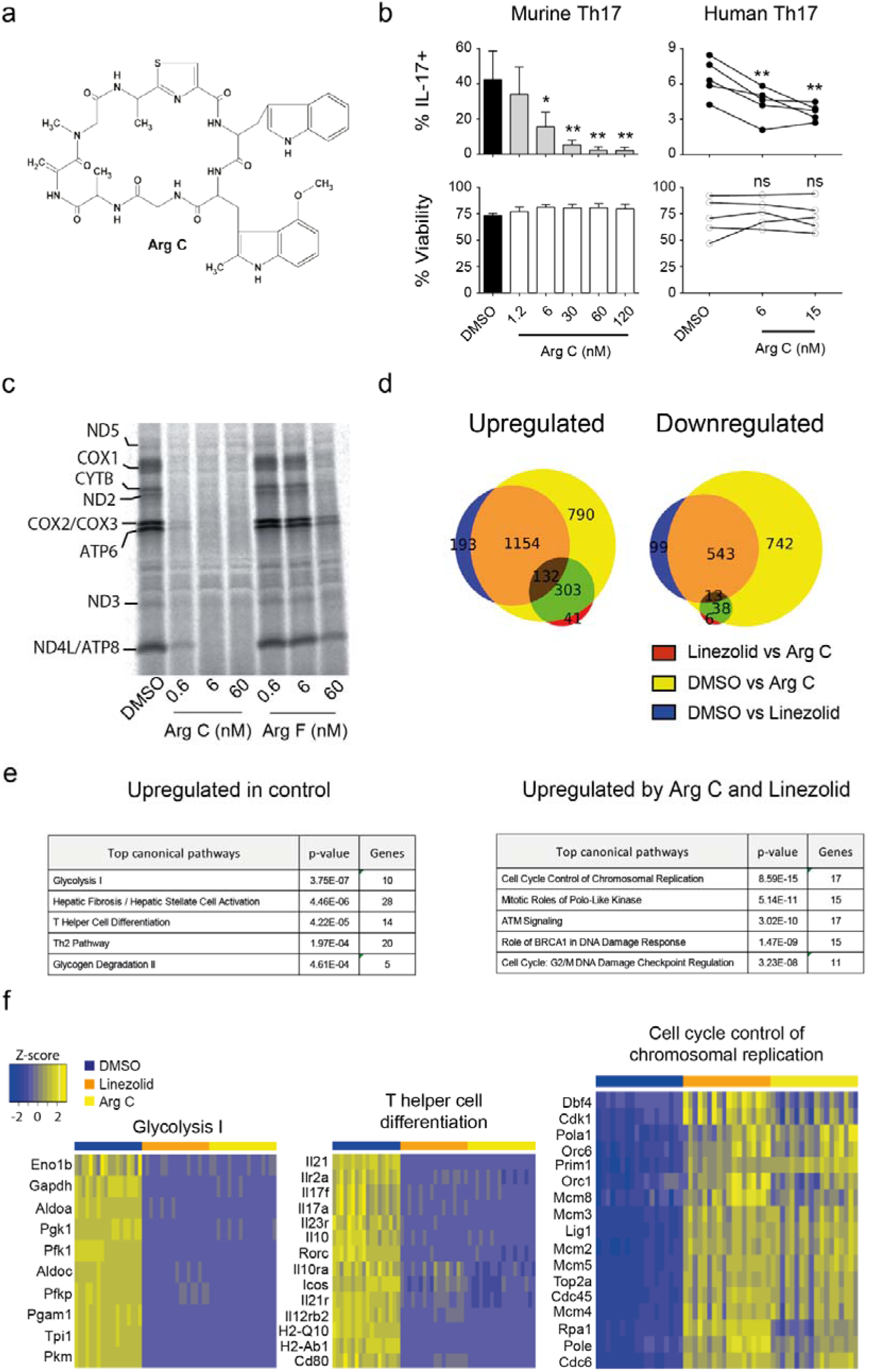
Inhibiting mitochondrial translation with Arg C inhibits Th17 cell function. **(a)** Chemical structure of Arg C. **(b, left)** Mouse naïve CD4^+^ T cells were cultured for 4d under Th17 polarizing conditions in the presence of Arg C (1.2, 6, 30, 60 and 120nM) or vehicle (DMSO). Cells were stained for intracellular IL-17A and gated on live CD4^+^ T cells. Bar graphs show % of IL-17A^+^ cells (top left) amongst total live CD4^+^ T cells (bottom left). **(b, right)** Human CD4^+^ CD45RO^−^ naïve T cells isolated from cord blood were cultured for 6d under Th17 polarizing conditions and treated with Arg C (6 and 15nM) or vehicle (DMSO). Cells were stained for intracellular IL-17A; gated on live CD4^+^ T cells. Graphs contain the data for each of 5 individual donors, indicated by dots connected by the same line. The values represent % of IL-17A^+^cells amongst live CD4^+^ cells (above) and the % of viable CD4^+^ cells (below). **(c)** Representative plot showing mitochondrial translation levels of EL-4 cells pre-incubated for 1h with Arg C (0.6, 6 and 60 nM), Arg F (0.6, 6 and 60nM) or vehicle (DMSO) and radioactively labeled with [^35^S]-methionine. **(d-f)** Mouse naïve CD4^+^ T cells were cultured under Th17 polarizing conditions in the presence of different concentrations of Arg C (60nM), linezolid (100µM) or DMSO. **(d)** Venn diagrams showing number of genes differentially up-(left) or downregulated (right) by linezolid vs Arg C, DMSO vs Arg C and DMSO vs linezolid. The overlap regions contain overlapping genes, found in both conditions **(e)** Tables show the top canonical pathways upregulated either by DMSO (left) or by both Arg C and linezolid (right) and **(f)** Heat maps showing up- or downregulation of genes among three selected relevant pathways. Results (**b left)** are the pooled means of technical replicates from three independent experiments. Representative results of two **(c)** experiments. Genes **(e)** are included if they were found to be up- or downregulated in both independent experiments performed. See also Figures S1 and S2.

Considering the marked differences in the activity of each compound, we focused on the most active metabolite, Arg C, for further studies (**Fig. 3a**). To assess whether Arg C exerts a general inhibitory effect on effector Th subset differentiation, we differentiated Th17, Th1 and Th2 under a broader range of Arg C concentrations. Addition of Arg C during mouse and human Th17 differentiation resulted in a dose dependent inhibition of the frequencies of IL-17A produced from these cells (**Fig. 3b**, above), without affecting their viability (**Fig. 3b**, below). Cytokine production by murine Th1 and Th2 cells were also affected, with IFN-γ and IL-13 production being equally reduced by Arg C (**Fig. S2a**). As expected, mEF-G1 inhibition by Arg C resulted in loss of mitochondrial translation, as measured by a near complete blockage of [^35^S]-methionine incorporation in mtDNA-encoded subunits in Arg C-treated EL-4 cells (**Fig. 3c**). A reduction of [^35^S]-methionine incorporation was also detectable after Arg F pretreatment (60nM). However, this effect was partial and only observed at a concentration 100 times higher than required for a similar inhibition with Arg C. This correlated with the lack of cytokine inhibition by Arg F at 60nM (**Fig S1b** and **c**).

Both linezolid and Arg C present as inhibitors of mitochondrial protein synthesis, despite their different molecular targets. To determine overall similarities between both drugs, we performed RNAseq analysis on Th17 cells exposed to linezolid, Arg C and DMSO. Differential gene expression analysis showed most genes significantly up- or downregulated by linezolid were equally up- or downregulated by Arg C after 96h (**Fig. 3d**). Among the top downregulated pathways by both linezolid and Arg C, were those corresponding to glycolysis and Th cell differentiation (**Fig. 3e, f**). Noteworthy, pathways significantly upregulated in linezolid and Arg C were associated with cell cycle control and response to DNA damage (**Fig. 3e, f**). Importantly, genes downregulated by both linezolid and Arg C in the “T helper cell differentiation” pathway included several Th17-associated genes, such as *Il17a*, *Il17f*, *Il23r, Rorc* and *Il21* (**Fig. 3f**). Taken together, our results demonstrate a high degree of similarity between the effects of linezolid and Arg C on Th17 cells.

### Proliferating T cells depend on functional mitoribosomes to maintain their ETC integrity and effector function

Naïve T cells have a lower proliferation rate, cellular respiration, and a decreased mitochondrial proteome when compared to activated T cells (39–42). T cell activation leads to several rounds of cell division, which requires the biogenesis of mitochondrial proteins. We therefore postulated that activated T cells would have a higher requirement for newly synthesized ETC subunits, including those encoded by mtDNA. To assess the reliance of differentiating and naïve T cells on mitochondrial translation, we measured the intracellular content of ND1 and COX1 throughout a culture with Arg C. After 48h of treatment, both ND1 and COX1 levels sharply decreased exclusively in Arg C-treated differentiating Th17 cells. Arg F, at sub-inhibitory concentrations (60nM), failed to reduce the levels of either subunit **(Fig. 4a**, left). Arg C, like linezolid, did not reduce the expression levels of the nuclear-encoded protein SDHA. Importantly, naïve T cells partially retained both COX1 and ND1 even after 120h of Arg C-treatment (**Fig. 4a**, right), showing that activated T cells are more sensitive to inhibition of this pathway than when maintained in a naïve state.

**Figure 4:**
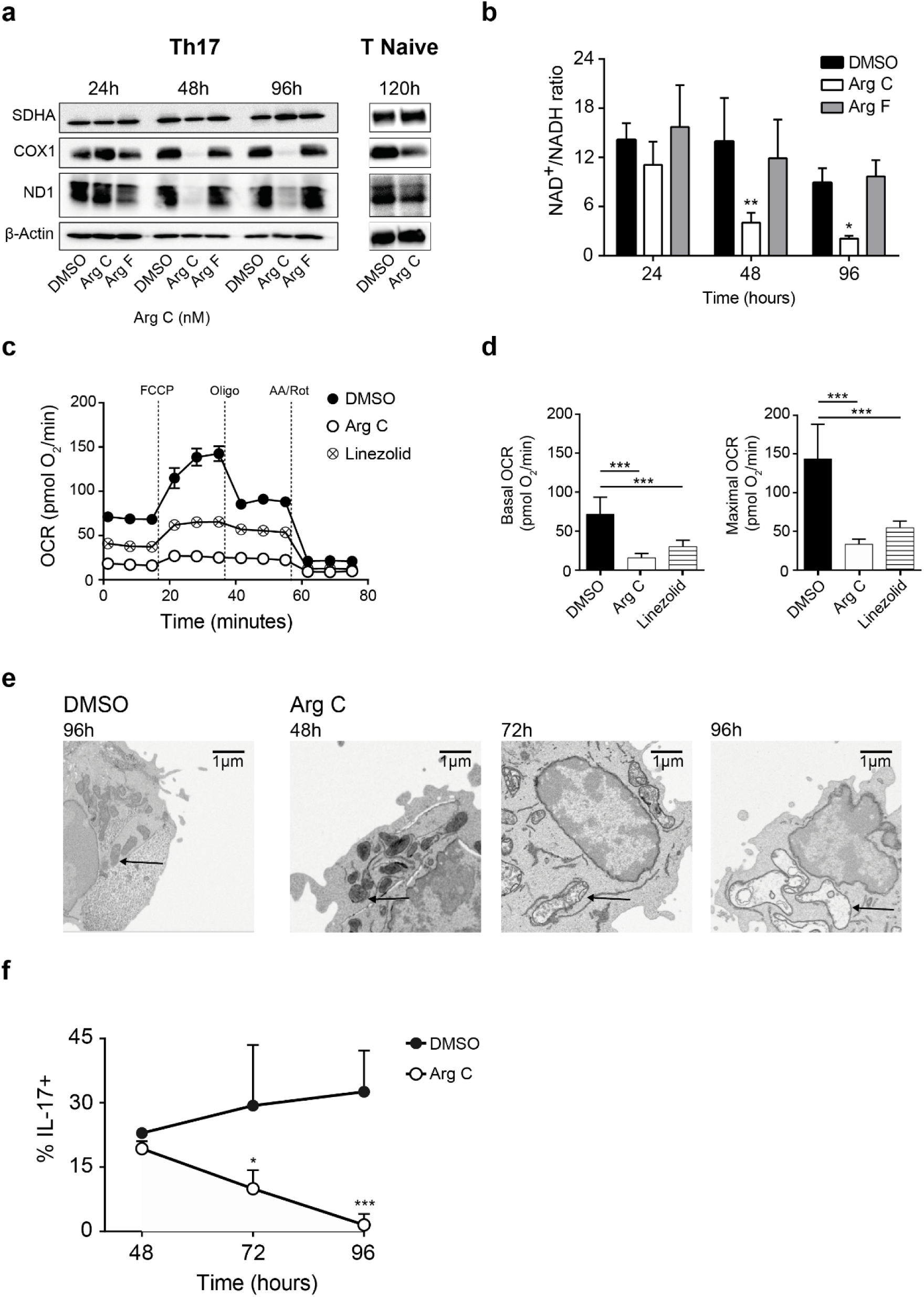
Arg C treatment selectively affects high proliferating cells, leading to mitochondrial dysfunction. **(a-f)** Naïve CD4^+^ T cells were cultured under Th17 polarizing conditions in the presence of Arg C (60nM), Arg F (60nM), linezolid (100µM) or vehicle (DMSO). **(a)** Whole cell lysates were analyzed for total protein levels of SDHA, ND1 and COX1 by western blot after 24, 48 and 96h of culture. **(b)** NAD^+^/NADH ratio measured after 24, 48 and 96h of culture. **(c)** Representative plot measuring real-time changes in oxygen consumption rate (OCR) at 72h of culture and **(d)** respective basal (before drug addition) and maximal (after FCCP addition) respiratory activity of the cells. OCR is reported as picomoles (pmol) of O_2_ per minute. Oligo, oligomycin; FCCP, carbonyl cyanide-p-trifluoromethox-yphenyl-hydrazon; Rot, rotenone; AA, antimycin A. **(e)** Representative electronic microscopy images show mitochondria morphology of Arg C-(60nM) or DMSO-treated naïve T cells from 48h of culture. **(f)** Cells were stained for intracellular IL-17A. Graph shows total % IL-17A^+^ cells amongst total live CD4^+^ T cells at 48, 72 and 96hr of culture. Statistical significance was determined using 2-way ANOVA with Bonferroni multiple corrections test, compared to vehicle-treated cells. Representative results of three **(a)**, five **(d)** and one **(e)** experiments are shown. Results are pooled means of technical replicates from three **(b, f)** and five **(d)** independent experiments. See also Figures S3 and S4.

Mitochondrial respiration has been described to support essential metabolic needs in proliferating cells through regeneration of electron accepting co-factors such as NAD^+^ or FAD using oxygen as the terminal electron acceptor (43). Consistent with the findings from Sullivan *et al* (43), the impact of both Arg C and linezolid on mitochondrial respiration resulted in a decreased intracellular NAD^+^/NADH ratio after 48h (**Fig. 4b** and **S3a**). A reduced NAD^+^/NADH ratio leads to allosteric inhibition of the enzyme complex pyruvate dehydrogenase, which converts pyruvate into acetyl-CoA by decarboxylation. In these conditions, we speculated that pyruvate would preferentially be converted into lactate (24, 44, 45). This hypothesis was corroborated by a drop in pH of the culture medium together with a slight increase in lactate production by Arg C-treated cells as compared to Arg F or DMSO (**Fig. S3b, c**). Deficient electron flow from the pyridine nucleotide NADH, to oxygen, together with a lack of mitochondrial-encoded subunits of complex V, F_1_F_o_-ATPase (namely ATP6 and ATP8), would hinder OXPHOS-derived ATP production. Accordingly, the ADP/ATP ratio was significantly increased in Arg C-treated cells compared to controls (**Fig. S3d**).

As expected, Arg C and linezolid-treated cells lost the ability to carry out mitochondrial respiration, illustrated by a lower basal and FCCP-stimulated maximal cellular respiration (**Fig. 4c, d**). Mitochondrial dysfunction by Arg C was also illustrated by electron microscopy imaging of differentiating Th17 cells. In contrast to the round shaped mitochondria characteristic of Th cells (40), Arg C treated cells presented elongated mitochondria with loosened cristae (**Fig. 4e**). After 96h of culture, this resulted in sac-shaped or swollen mitochondria, comparable to the morphology of this organelle in cells devoid of mitochondrial DNA (ρ^0^ cells) (46). Similar to previous reports on ρ^0^ cells (47), Arg C-treated cells were able to maintain a mitochondrial membrane potential (**Fig. S3e**). Importantly, while loss of mtDNA-encoded subunits was evident at 48h, IL-17 production was only reduced at 72h (**Fig. 4f**). This suggested that initial differentiation steps were unaffected, as evidenced by the capacity for Arg C-treated cells to produce IL-17 until 48h. To confirm this, we measured the expression of the Th17-cell master transcription factor RORγt. Indeed, Arg C did not affect expression of the transcription factors T-bet, GATA-3 or RORγt in Th1, Th2 and Th17 cells, respectively (**Fig. S4a-c**). This indicates that Arg C impairs cytokine production and not the initial Th cell transcriptional program.

Considering our transcriptomic data suggested cell cycle pathways to be dysregulated (**Fig. 3e, f**) we investigated the relationship between proliferation and loss of cytokine production caused by Arg C. Using cells labelled with CellTrace^TM^ we observed that proliferation under Arg C treatment was mildly impaired toward the final stages of differentiation (**Fig. S4d**). Importantly, this moderate decline in cellular proliferation was not the main cause of cytokine inhibition, as all cells from across each individual cycle showed significantly reduced IL-17 production after 72h of culture (**Fig. S4e**). Therefore, Arg C’s inhibitory effect on cytokine production is not merely a consequence of impaired cellular division of effector cells.

### *Gfm1* loss results in deficient mitochondrial translation and impaired Th17 effector function

Aside from targeting mitochondrial ribosomes, linezolid is also a weak inhibitor of monoamine oxidase (48). In addition, the Argyrins have also been described as inhibitors of the proteasome (49, 50). To exclude that any of these alternative side effects were responsible for the immunosuppressive effects seen by these drugs, we genetically targeted mitochondrial protein synthesis in T cells. As the target of linezolid is the mitoribosome, which is encoded fully by mtDNA, we focused on *Gfm1*, the gene encoding mEF-G1, located on chromosome 3 (51). Given that ubiquitous deletion of *Gfm1* is lethal at pre-weaning stages, we generated an inducible T cell specific *Gfm1* deficient mouse strain (T-Gfm1Δ). Mice expressing a CD4-specific, tamoxifen-inducible Cre enzyme (Cd4Cre^ERT2/wt^) were crossed with mice carrying a loxP-flanked exon 4 in the *Gfm1* gene (Gfm1^flox/flox^) (**Fig. 5a**). In contrast to complete *Gfm1* knockout mice, T-Gfm1Δ mice are viable and express normal levels of COX1 in their CD4^+^ T cells, indicative of functional mitochondrial translation (**Fig. 5b and S5a**). Upon treatment with 4-hydroxytamoxifen (4-OHT), COX1 expression levels significantly dropped in CD4^+^ T cells from T-Gfm1Δ mice in comparison with those still harboring *Gfm1* WT alleles, either in a haplo-sufficient form (Cd4Cre^ERT2/wt^Gfm1^flox/wt^) (**Fig. 5b)** or lacking Cre-recombinase activity (Gfm1^flox/flox^Cd4Cre^wt/wt^) (**Fig. S5a**). Similar to Arg C, differentiation of 4-OHT-treated naïve T cells isolated from T-Gfm1Δ mice under Th17-skewing conditions yielded a significantly reduced frequency of IL-17A-expressing cells (**Fig. 5c, Fig. S5b**). Finally, we demonstrate that tamoxifen-mediated genetic deletion of *Gfm1* in CD4^+^ cells in T-Gfm1Δ mice, leads to significant protection from the development of EAE. Mice retaining a functional *Gfm1* allele in CD4^+^ T cells (Gfm1^flox/wt^) are more susceptible to symptoms of the disease (**Fig. 5d)**. Taken together, our results indicate that loss of mitochondrial translation in T cells, either genetically or pharmacologically, restrains Th17 function. We further show that mitochondrial translation is necessary for T cell pathogenicity *in vivo* and could represent a therapeutic target to halt Th cell aberrant responses.

**Figure 5:**
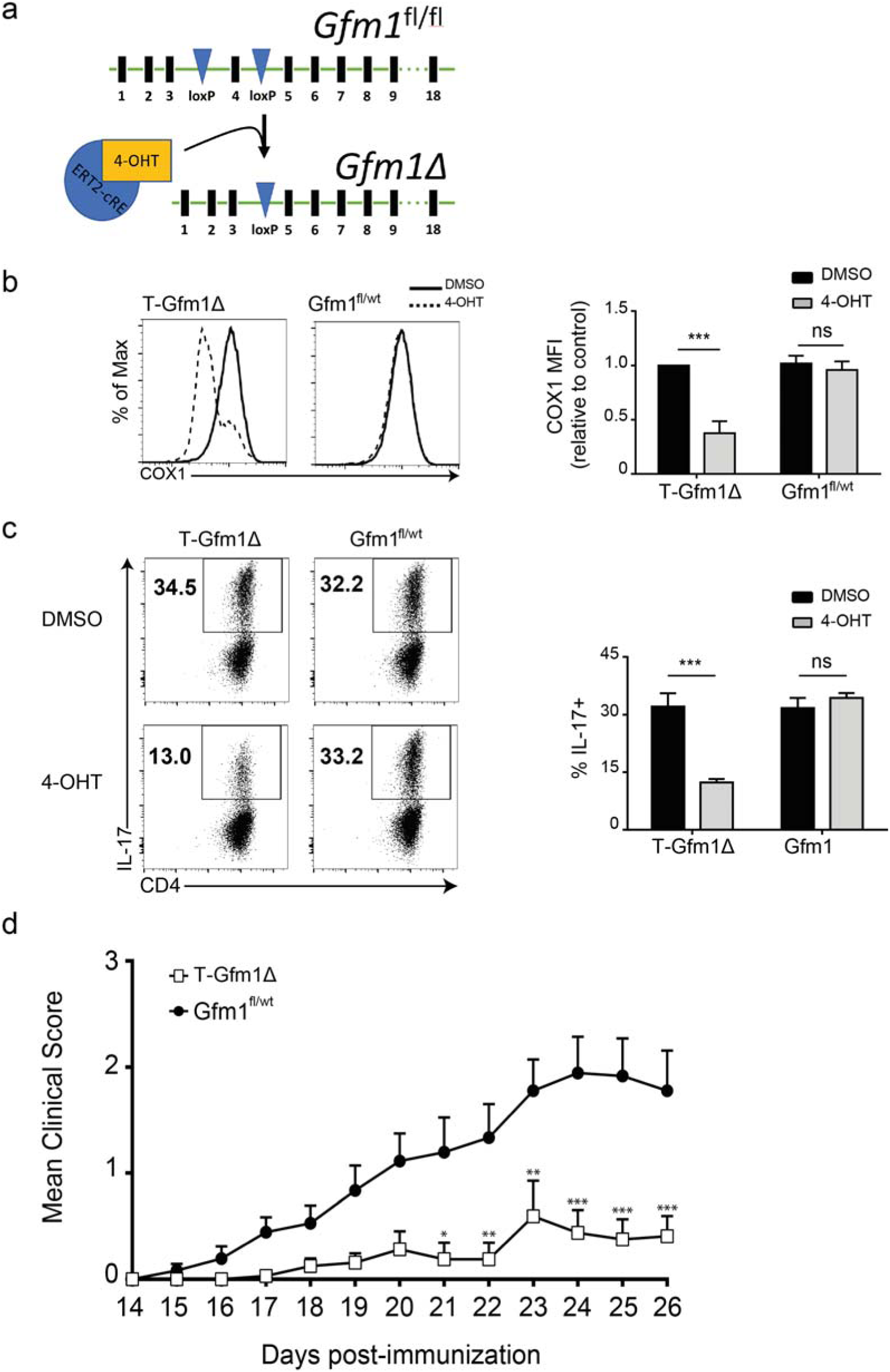
T cell-specific *Gfm1* deletion attenuates Th17 cytokine production and mitigates development of EAE in MOG-immunized mice. **(a)** Graphical image highlighting the generation of CD4-specific, tamoxifen-inducible Cre mouse line, which deletes the loxP-flanked exon 4 of the *Gfm1* gene (Cd4Cre^ERT2/wt^ Gfm1^flox/flox^) after exposure to tamoxifen. **(b,c)** Sorted naïve CD4^+^ cells from T-Gfm1Δ mice (Gfm1^flox/flox^Cd4Cre^ERT2/wt^) and littermate haplosufficient controls (Gfm1^flox/wt^Cd4Cre^ERT2/wt^) were pre-treated with 300nM tamoxifen (4-OHT) or DMSO and maintained in a naïve state 12 days, prior to being differentiated under Th17 skewing conditions for 4 days. Intracellular levels of COX1 and IL-17A were quantified by FACS. **(b)** Representative histograms (left) show COX1 expression amongst CD4^+^ T cells. Bar graph (right) shows COX1 median fluorescence intensity (MFI) relative to DMSO control. **(c)** Representative dot plots (left) show percentage of IL-17A^+^ amongst live CD4^+^ T cells. Bar graph (right) shows pooled means of the percentage of IL-17A^+^ CD4^+^ T cells. **(d)** T-Gfm1Δ mice (Gfm1^flox/flox^Cd4Cre^ERT2/wt^) and haplosufficient controls (Gfm1^flox/wt^Cd4Cre^ERT2/wt^) were immunized with MOG_35-55_ in CFA and pertussis toxin to induce EAE. Tamoxifen was administered once to every mouse, 3 days after induction of EAE. The graph contains the mean clinical score for T-Gfm1Δ mice (Gfm1^flox/flox^Cd4Cre^ERT2/wt^) and haplosufficient controls (Gfm1^flox/wt^Cd4Cre^ERT2/wt^). Bar graphs (**b, c**) contain the pooled means from the technical replicates of three individual experiments. Plots are obtained from the pooled data of two **(d)** individual experiments, with error bars showing the S.E.M of the pooled scores. Statistical significance was determined using 2-way ANOVA with Bonferroni multiple corrections test compared to “untreated” mice. See also Figure S5.

## Discussion

It is well established that some RAbos can inhibit mammalian mitochondrial ribosomes. Furthermore, certain RAbos were previously described as immunosuppressive. For example, chloramphenicol diminishes neutrophil extracellular trap release and doxycycline inhibits B cell class switching and IgM secretion (52, 53). However, how the reduction of immune cell function occurred was mechanistically not well understood. We show here that inhibition of mitochondrial protein synthesis by antibiotics can reduce Th cell cytokine secretion. Certain clinical observations suggest that linezolid is immunosuppressive, even though this has never been officially proposed nor investigated. As previously mentioned, linezolid treatment is known to confer increased risk of fungal and gram-negative infections (14, 15). Th17 cell function is necessary in mounting antifungal and antibacterial responses. Linezolid is active exclusively against gram-positive bacteria, and we postulate that linezolid facilitates opportunistic fungal or gram-negative infections by dampening Th17 function. Importantly, we found linezolid to inhibit both mitochondrial translation and Th17 cell function *in vitro* at concentrations routinely achieved during clinical administration (54). We show in a T cell-mediated autoimmunity model of multiple sclerosis (EAE) that linezolid prevents the development of self-reactive T cells. Thus, repurposing linezolid as an immunosuppressive drug resulted in significant protection from pathogenic T cell infiltration and led to absent (or mild) EAE symptoms, reflected by lower clinical scores. However, future studies will be necessary to determine if T cell-suppression by linezolid contributes to the progression of infections by gram-negative or fungal pathogens.

To further study the role of mitochondrial translation in T cells, we searched for alternative methods to inhibit this process. In light of the indispensable role of mEF-G1 in catalyzing the translocation of mitoribosomes during mitochondrial protein translation (37, 55), we made use of Arg C to hinder mEF-G1’s activity during Th differentiation. Murine mEF-G1 shares 89% sequence identity with the human homologue mEF-G1 (51), and accordingly, Arg C was active against both human and mouse Th17 cells. The similarities in the gene transcriptional changes of Arg C- and linezolid-treated cells further confirm that targeting mitochondrial translation with either compound leads to analogous cellular effects. Using Arg C, we were able to determine that differentiating T cells are highly reliant on a functional mitochondrial protein synthesis machinery to keep up with ETC biosynthetic demands. Notably, although Arg C inhibited mitochondrial protein synthesis after 1h of treatment, the lack of mitochondrial-encoded and -translated ETC subunits was first evidenced after 48h of cell activation. At 24h the majority of cells have not yet proliferated (56) and, presumably, did not need to synthesize large amounts of new ETC subunits. Between 24-48h, cell growth and division would require an increase in mitochondrial mass and proteome. Therefore, an impaired mitochondrial translation machinery would render cells unable to keep up with the cellular demand for an increased synthesis of ETC subunits. Ultimately, this would “dilute” pre-existing mitochondrial-encoded proteins such as COX1 and ND1 amongst daughter cells. Consequently, interfering with mitochondrial translation might be particularly detrimental for rapidly dividing cells such as activated T cells (57), while non-proliferative cells should be able to retain their ETC subunits for longer. Accordingly, we show that naïve T cells exposed to Arg C for as long as 120h still expressed both COX1 and ND1.

Activated CD4^+^ T cells in the presence of Arg C are subjected to a gradual loss of ETC integrity which leads to reduced oxygen consumption and a concomitant loss of electron acceptors such as NAD^+^. To circumvent this, the cell is forced to convert pyruvate into lactate. As fermentation of glucose into lactate does not yield net NAD^+^ (39), over time the cofactor levels required to support essential oxidative steps in proliferating cells decrease. Accordingly, Th17 cells treated with Arg C display both a reduced NAD^+^/NADH ratio, respiration and increased lactate secretion. Interestingly, inhibition of mitochondrial translation by Arg C did not influence mitochondrial membrane potential. In this respect, studies on cells lacking mtDNA, termed ρ°, revealed that these cells display a mitochondrial membrane potential high enough to sustain the import of nuclear-encoded proteins. The mechanism is thought to involve an electrogenic exchange of ATP^4-^ for ADP^3-^ by the adenine nucleotide carrier and maintained by an incomplete F_1_F_o_-ATPase (47, 58, 59). This process would consume glycolysis-derived ATP and is in agreement with the strong increase of ADP/ATP ratio measured in Arg C-treated cells.

Upon activation, CD4^+^ T cells cycle must undergo several rounds of proliferation to develop into cytokine producing Th effector cells (60). While loss of COX1 and ND1 occurs at 48h, neither proliferation nor IL-17 production are affected at this time point. Instead, we observe that if mitochondrial translation remains inhibited, cells progressively become unable to produce IL-17. The down regulation of cytokines becomes evident from 72h onwards, present simultaneously in cells from all individual proliferation cycles. This highlights that Arg C’s cytokine inhibition and anti-proliferative phenotypes are uncoupled, and inhibition of proliferation *per se* cannot explain loss of IL-17 production. Arg C appears to strictly affect effector function, and not initial Th cell transcriptional programs. This is consistent with Arg C-treated Th1, Th2 and Th17 cells retaining their expression of master transcription factors T-bet, GATA-3 and ROR*γ*t, respectively.

While our results might suggest the repurposing of antibiotics as immunomodulatory drugs, this might be a double-edged sword given a potential negative impact on gut flora (25, 61). Of greater consequence, widespread usage of linezolid in an autoimmune setting might speed up the dissemination of resistant bacterial strains to this reserve antibiotic. Alternatively, antibiotics which are not routinely used in human medicine, such as thiamphenicol, could be further explored as immunomodulators. Importantly, the Argyrins display poor antibacterial and antifungal properties and are not currently used as antibiotics in the clinic (62). Furthermore, inhibition of mEF-G1 by Arg C blocked mitochondrial translation in Th17 cells at concentrations 4-5 orders of magnitude lower than linezolid. Even though we lacked the means to develop a good *in vivo* administration method for Arg C, we propose that further steps should be taken to adapt novel and more potent mitochondrial translation inhibitors for therapeutic use. To encourage such efforts, we investigated the role of mEF-G1 in Th17 cell function by generating an inducible T cell specific *Gfm1* deficient mouse strain. Tamoxifen-mediated deletion validated that mEF-G1 is necessary for T cells to maintain their effector function *in vitro*. Moreover, specifically deleting *Gfm1* in CD4^+^ cells at the onset period of EAE induction protected against disease symptoms and provided strong evidence that this pathway is necessary *in vivo*, for T cell-mediated pathogenicity. Taken together, our results provide strong evidence to consider the inhibition of mitochondrial translation as a novel approach to halt pathological Th cell responses. Additionally, we identify mEF-G1 as a promising target to control this cellular process and achieve immunomodulation and highlight the ability of RAbos to alter host cell functions.

## Acknowledgements

We thank all members of the Institute of Infection Immunology at TWINCORE for discussion and support. We thank Luciana Berod, Institute of Infection Immunology at TWINCORE, for critical reading and discussion of the manuscript. We would like to acknowledge the assistance of the Cell Sorting Core Facility of the Hannover Medical School supported in part by the Braukmann-Wittenberg-Herz-Stiftung and the Deutsche Forschungsgemeinschaft. We thank Heinrich Steinmetz from Helmholtz Centre for Infection Research for providing Arg C, Peter Rehling from University Medical Centre (Göttingen) for providing resources to conduct mitochondrial translation assays and the Department of Prenatal Medicine and Midwifery of the Medical School Hannover (MHH) for providing human cord blood. We would also like to thank the Teichmann Lab at the Wellcome Trust Sanger Institute and Dora Pedroso at Instituto de Ciência (IGC) for technical support. This work was supported by grants from the Deutsche Forschungsgemeinschaft CRC156 to T.S.; A.D.L. was supported by the Hannover School for Biomedical Drug Research (HSBDR) and L.A. has received funding from the European Union’s Horizon 2020 research and innovation program under the Marie Skłodowska-Curie grant agreement No 675395. B.R.B was supported by the National Institutes of Health R01 HL56067.

## Author Contributions

Conceptualization T.S.; Investigation, L.A., A.D.L., C.N.C., G.M.C., M.G., C.H., N.A., S.D., and S.L.; Resources, B.N.L., L.S., M.K., and R.M.; Writing and Visualization, L.A., A.D.L. C.N.C., and T.S., Critical Revision, L.F.M., B.N.L., B.R.B.; Supervision, Project Administration and Funding Acquisition, T.S.

## Author Information

The authors declare no competing financial interests. Correspondence and requests for materials should be addressed to T.S. (Sparwasser).

## Methods

### Mice

C57BL/6 mice were purchased from Jackson Laboratories. Gfm1^flox/flox^ were generated by crossing mice carrying the heterozygous Gfm1^tm1a(EUCOMM)Wtsi^ “knockout first” allele (purchased from EMMA mouse repository) with Flp-expressing mice to obtain the conditional allele (63). T-Gfm1Δ were generated by crossing Cd4Cre^ERT2/wt^ (64) to Gfm1^flox/flox^. Sex- and age-matched mice between 7 and 20 weeks of age were used for all experiments. All mice were bred and maintained under specific pathogen free conditions at the animal facility of the Helmholtz Centre for Infection Research (HZI, Braunschweig, Germany) or at TWINCORE (Hannover, Germany). All animal experiments were performed in compliance with the German animal protection law (TierSchG BGBl. I S. 1105; 25.05.1998) and were approved by the Lower Saxony Committee on the Ethics of Animal Experiments as well as the responsible state office (Lower Saxony State Office of Consumer Protection and Food Safety under the permit number 33.9-42502-04-12/0839).

### Mouse T cell cultures

CD4^+^ CD25^−^ T cells were isolated *ex vivo* from spleens and lymph nodes of mice by enrichment with EasyStep^TM^ Mouse CD4+ Isolation Kit (Stemcell Technologies) in combination with a biotin-conjugated antibody against CD25, followed by immunomagnetic negative isolation. The purity of the isolated cells was ∼90%. RPMI 1640 GlutaMAX^TM^ medium or IMDM GlutaMAX^TM^ medium (both from Life Technologies) was used for Th1, Th2 and Treg or Th17 cultures, respectively. Medium was supplemented with 10% heat-inactivated FCS (Biochrom), 500 U penicillin-streptomycin (PAA laboratories) and 50 μM β-mercaptoethanol (Life Technologies). For Th17 induction, 2 to 3×10^5^ naïve T cells were cultured for 4 days with plate-bound αCD3ε (10 μg mL^−1^, clone 145-2C11; Bio X Cell), αCD28 (1 μg/mL, clone 37.51; Bio X Cell), αIFN-γ (5 μg/mL, clone XMG1.2; Bio X Cell), αIL-4 (5 μg mL^−1^, clone 11B11; Bio X Cell), rhTGF-β1 (2 ng mL^−1^; Peprotech), rmIL-6 (7.5 ng mL^−1^; Peprotech) and rmIL-1β (50 ng mL^−1^; Peprotech). For Th1 induction, 1×10^5^ naïve T cells were cultured for 4 days in the presence of plate-bound αCD3ε (10 μg mL^−1^), αCD28 (1 μg mL^−1^), rmIL-12 (50 ng mL^−1^) and αIL-4 (5 μg mL^−1^). For Th2 induction 1×10^5^ naïve T cells were cultured for 4 days with plate-bound αCD3ε (10 μg mL^−1^, clone 145-2C11; Bio X Cell) and αCD28 (10 μg mL^−1^, clone 37.51; Bio X Cell), αIFN-γ (10 μg mL^−1^, clone XMG1.2; Bio X Cell), αIL-12 (10 μg mL^−1^, clone 17.8; Bio X Cell) and rmIL-4 (1 μg mL^−1^, Preprotech). For Treg induction, 2.5×10^4^ naïve T cells were cultured for 4 days in the presence of plate-bound αCD3ε (5 μg mL^−1^), αCD28 (1 μg mL^−1^), rhIL-2 (200 U mL^−1^; Roche Applied Science) and rhTGF-β1 (1 ng mL^−1^). On day 2, rhIL-2 (200 U mL^−1^) was added again. Arg C, Arg F, linezolid (Chem-Impex), tigecycline hydrate (Sigma) and thiamphenicol (Chem-Impex) were added at the indicated concentrations at the onset of the culture. For proliferation analysis, naïve T cells were labeled using 5µM CellTrace™ Violet Cell Proliferation Kit (Life Technologies) after magnetic sorting.

### Human T cell cultures

Human cord blood samples were obtained from the Department of Prenatal Medicine and Midwifery of the Medical School Hannover (MHH). All work with human blood samples was approved by the local ethics committee and informed consent was obtained from all subjects. After Ficoll (Biocoll) gradient, naïve CD44^+^ T cells were enriched by magnetic separation using the EasySep™ Human Naïve CD4+ T Cell Isolation Kit (Stemcell Technologies). 55×10^4^ T cells were cultured for 6 days in the presence of plate-bound αCD3ε (5 μg mL^−1^), in X-Vivo 15 medium (Lonza), supplemented with 2% heat-inactivated FCS (Biochrom), 500 U penicillin-streptomycin (PAA laboratories) and 50 μM β-mercaptoethanol (Life Technologies). To polarize the cells towards a Th17 phenotype, the medium was supplemented further with rhIL-1β (10 ng mL^−1^, R&D Systems), rhIL-23 (20 ng mL^−1^, R&D Systems), rhIL-6 (20 ng mL^−1^; Peprotech), rhIL-21 (20 ng mL^−1^; Peprotech), rhTGF-β1 (3 ng mL^−1^; Peprotech), αCD28 (500ng mL^−1^) and 20mM NaCl. Arg C and linezolid were added at the indicated concentrations from day 0.

### Naïve T cell maintenance and *Gfm1* deletion

Naïve T cells (CD4^+^ CD25^−^ CD62L+ CD44^low^) were sorted from spleens and lymph nodes of mice and maintained in RPMI 1640 GlutaMAX^TM^ medium (from Life Technologies) supplemented with 10% heat-inactivated FCS (Biochrom), 1X Antibiotic Antimycotic Solution (Sigma) and 50 μM β-mercaptoethanol (Life Technologies), containing rmIL-7 (20ng mL^−1^, Peprotech). rmIL-7 (10ng mL^−1^) was re-added at the 3^rd^, 5^th^, 7^th^, 9^th^ and 11^th^ days of culture. To induce tamoxifen-mediated deletion *in vitro*, (Z)-4-Hydroxytamoxifen (Sigma) at 300nM was added at the beginning of the culture. After 12 days, naïve T cells were harvested, counted, and re-plated at a density of 1.5×10^5^ cells per well under Th17-polarizing conditions as described above. For *in vivo* deletion of *Gfm1* during EAE induction, mice were administered with a single dose of 2mg of Tamoxifen (Santa Cruz) dissolved in corn oil intraperitoneally, 3 days after EAE induction.

### Flow cytometry

Monoclonal antibodies specific against the following mouse antigens were purchased from Affymetrix/eBioscience: CD4 (GK1.5), IL-17A (eBio17B7), IL-13 (eBio13A), IFN-γ (XMG1.2), T-bet (eBio4B10), RORγt (B2D), GATA-3 (TWAJ), FoxP3 (FJK-16s), CD154 (MR1). COX1 AF488 was purchased from Abcam. For human cells, the following antibodies purchased from Affymetrix/eBioscience were used: CD4 (SK3), IL-17A (eBio64DEC17). For analysis of surface markers, cells were stained in PBS containing 0.25% bovine serum albumin (BSA) and 0.02% azide. Dead cells were excluded by LIVE/DEAD® Fixable Dead Cell Stain Kit (Life Technologies). For mouse intracellular cytokine staining, cells were stimulated with phorbol 12-myristate 13-acetate (PMA) (0.1 μg mL^−1^; Sigma-Aldrich) and ionomycin (1 μg mL^−1^; Sigma-Aldrich) for 2 h followed by brefeldin A (5 μg mL^−1^) for 2 h and fixed using 2% PFA fixation followed by permeabilization with PBS containing 0,25% BSA and 0,5% of saponin. Human intracellular staining was done as described above, but brefeldin A was added simultaneously with PMA and ionomycin instead. For transcription factor staining, cells were instead fixed with Foxp3/Transcription Factor Fixation/Permeabilization Kit (Affymetrix/eBioscience) according to manufacturer’s instructions prior permeabilization. Stained cells were acquired on a CyAn^TM^ ADP (Beckman Coulter) or a LSR II (Becton Dickinson) and data was analyzed with FlowJo software (Tree Star, Inc.).

### Western blot

Whole cell lysates were prepared using lysis buffer (Pierce™ RIPA buffer, Thermo Scientific) supplemented with complete EASYpack Mini Protease Inhibitor Cocktail and PhosSTOP Phosphatase Inhibitor (both from Roche Applied Science). Cell lysates were separated by SDS-gel electrophoresis and transferred to PVDF membranes (Merck Millipore). Immunoblotting was performed using the following antibodies from Cell signaling: mouse anti-β-actin, goat-anti-rabbit (Horseradish peroxidase conjugated) and anti-mouse (Horseradish peroxidase conjugated). The following antibodies were purchased from Abcam: mouse anti-Sdha, rabbit anti-ND1, mouse anti-COX1.

### Metabolism analysis

Naïve CD4^+^ T cells were plated at a density of 1×10^5^ cells per well under Th17-polarizing conditions containing the indicated drug, harvested after 72 hours of culture and plated on 96-well XF cell culture microplates in XF assay medium (pH 7.4, both from Agilent) supplemented with D-glucose (20mM) and L-glutamine (2mM, Gibco) with a density of 3×10^5^ cells per well. Microplates were incubated for 30 min at 37 °C in a non-CO_2_ incubator and subjected to real-time analysis of OCR using an XF96 Extracellular Flux Analyzer (Agilent). For the mitochondrial stress assay analysis, the XF Mitochondrial Stress Test was performed according to manufacturer’s instructions, using subsequent injections of FCCP (0.8 μM), oligomycin (1 μM), and rotenone and antimycin A (0.5μM).

### Mitochondrial membrane potential measurement

Cells were stained at the indicated time points with tetramethylrodamine, ethyl ester (TMRE, Abcam®) according to manufacturer’s instructions. After 20 min incubation with TMRE at 37°C, cells were washed and TMRE was detected by flow cytometry.

### Lactate measurement

Lactate concentration in the extracellular medium of T cells cultured under Th17-polarizing conditions for 96h was performed using the Lactate Assay kit from Sigma-Aldrich according to manufacturer’s instructions.

### NAD^+^/NADH ratio measurement

NAD^+^/NADH ratio were measured in T cells cultured under Th17-polarizing conditions for 24, 48 and 96h using NAD/NADH-Glo^TM^ Assay (Promega) according to manufacturer’s instructions.

### ADP/ATP ratio measurement

ADP and ATP levels were measured in T cells cultured under Th17-polarizing conditions for 96h using the ADP/ATP Ratio Assay kit (Abnova) according to manufacturer’s instructions.

### Mitochondrial translation assay (or [^35^S]-methionine incorporation assay)

EL-4 cells were treated for 1h with the indicated compounds or vehicle and starved for 20 min before cytosolic translation blocking. [^35^S]-Methionine was added to the culture and incubated for 1h. Cells were subsequently harvested, lysed and subjected to electrophoretic separation in an SDS-gel. Mitochondrial translation products were detected by autoradiography.

### Electron microscopy analysis

Samples were incubated in freshly prepared fixative (2% paraformaldehyde (PFA, EMS), 2.5% gluteraldehyde (GA, EMS) in 0.1M Sodium Cacodylate (EMS) buffer, pH 7.4) at RT for 30 minutes. Fixative was removed by washing 5 × 3 minutes in 0.1M cacodylate buffer. 100µl cell suspension was embedded by centrifugation (5’ at 1230 g at 30°C) in 100µl 2% Low Melting Agarose (LMA, Gibco BRL) in 0.1M cacodylate buffer. Samples were then incubated in 2% osmium tetroxide (OsO4, EMS), 1.5% potassium ferricyanide (EMS) in 0.1M cacodylate buffer for 60 minutes at RT. After washing in H2O samples were incubated in 1% Thiocarbohydrazide (TCH, EMS) in water for 20 minutes. TCH was removed by washing 5 × 3 minutes in water. Then, a second incubation in OsO4 (2% OsO4 in H2O) for 30 minutes at RT. After washing in H2O for 5 × 3 minutes, samples were incubated overnight at 4°C in Uranyl Acetate Replacement (UA, EMS) in H2O (1:3). The next day, UAR was removed by washing in H2O for 5 × 3 minutes. The cells were then incubated for 30 minutes in Walton’s lead solution at 60°C. After final washing steps the samples were dehydrated using increasing EtOH concentration (70%, 90%, 2x 100%), for 5 minutes each followed by 2 × 10 minutes of propylene oxide (Aldrich). Subsequent infiltration with resin (Spurr, EMS) was done by first incubating in 50% resin in propylene oxide for 2 hours, followed by at least 3 changes of fresh 100% resin (including 1 overnight incubation). Next, samples were embedded in fresh resin and cured in the oven at 65 °C for 72 hours. For FIB-SEM imaging, embedded cells were mounted on aluminium SEM stubs (diameter 12 mm) and samples were coated with ∼8nm of Platinum (Quorum Q150T ES). FIB-SEM imaging was performed using a Zeiss Crossbeam 540 system with Atlas5 software. The Focussed Ion Beam (FIB) was set to remove 5nm sections by propelling Gallium ions at the surface. Imaging was done at 1.5kV using an ESB (back-scattered electron) detector.

### Dataset, quality control and RNA-seq analysis

The Raw FASTQ files were quality controlled, trimmed and mapped to the Mus musculus reference genome (GRCm38.p5) using the RNA-seq Analysis workflow of QIAGEN’s CLC Genomics Workbench 8. Expression levels were estimated as RPKM (Reads Per Kilobase of exon model per Million mapped reads) (65). Exploratory analysis revealed gene expression patterns following treatment of naïve T cells labelled with CellTrace™ Violet Cell Proliferation Kit and cultured with DMSO, linezolid and Arg C for 96h. Prior to RNA isolation, live CD4+ cells were sorted from each individual proliferation cycle. Differential expression analysis was performed between the treatment groups. The statistical significance threshold level of 0.05 with Bonferroni corrected FDR p-value and absolute fold change of 2 was used as a cutoff for differentially expressed genes. We compared the number of overlapping differentially expressed genes from the three treatment groups at each time points using a proportional Venn-diagram plotted using the Python matplotlib_venn library. Heatmap visualization for expression dynamics for the selected genes were plotted using the ComplexHeatmap R package. Pathway analysis was performed by uploading the processed RNA-seq data to Ingenuity Pathway Analysis (IPA, QIAGEN).

### Experimental autoimmune encephalomyelitis (EAE)

EAE was induced by subcutaneous immunization with 200 μg MOG35–55 peptide (HZI, Braunschweig) emulsified in CFA (Sigma-Aldrich), followed by intravenous injection of 200 ng pertussis toxin (Sigma-Aldrich) on days 0 and 2. To quantify disease severity, scores were assigned daily on a scale of 0–5 as follows: 0-no paralysis, 1-limp tail, 2-limp tail and partial hind leg paralysis, 3-complete hind leg paralysis, 4-tetraparesis and 5-moribund or dead. Animals were euthanized if score reached grade 3.5 for two consecutive days and subsequently scored as 5 the following days. To determine CNS infiltrates, mice were sacrificed at day 15 or 16 post-induction and cell suspensions from brain and spinal cord were prepared, as described previously (66). Single suspension of cells were cultured in 48-well plates in the presence of 30 μg mL^−1^ MOG35–55 for 2h followed by addition of Brefeldin A (5 μg mL^−1^; Affymetrix/eBioscience) for 4h. Finally, cells were analyzed by FACS for expression of surface markers and cytokine production. Where indicated, linezolid or vancomycin were administered daily after day 3 post-induction by oral gavage (4mg per mouse). Additionally, mice had *ad libidum* access to drinking water supplemented with 1mg mL^−1^ of the indicated antibiotic.

### Statistical analysis

Data analysis was performed using GraphPad Prism Software. Unless otherwise stated, statistics were calculated using One-way ANOVA with Bonferroni multiple comparisons test. Means are given as ±s.d. or where indicated as ±s.e.m. with *p*-values considered significant as follows: *\*p < 0.05; **p < 0.01* and *\*\*\*p < 0.001*. When appropriate (data originating from different human donors), the values for different treatments were matched and corrected with the Geisser-Greenhouse method.

## Figure Legends

**Figure S1.**
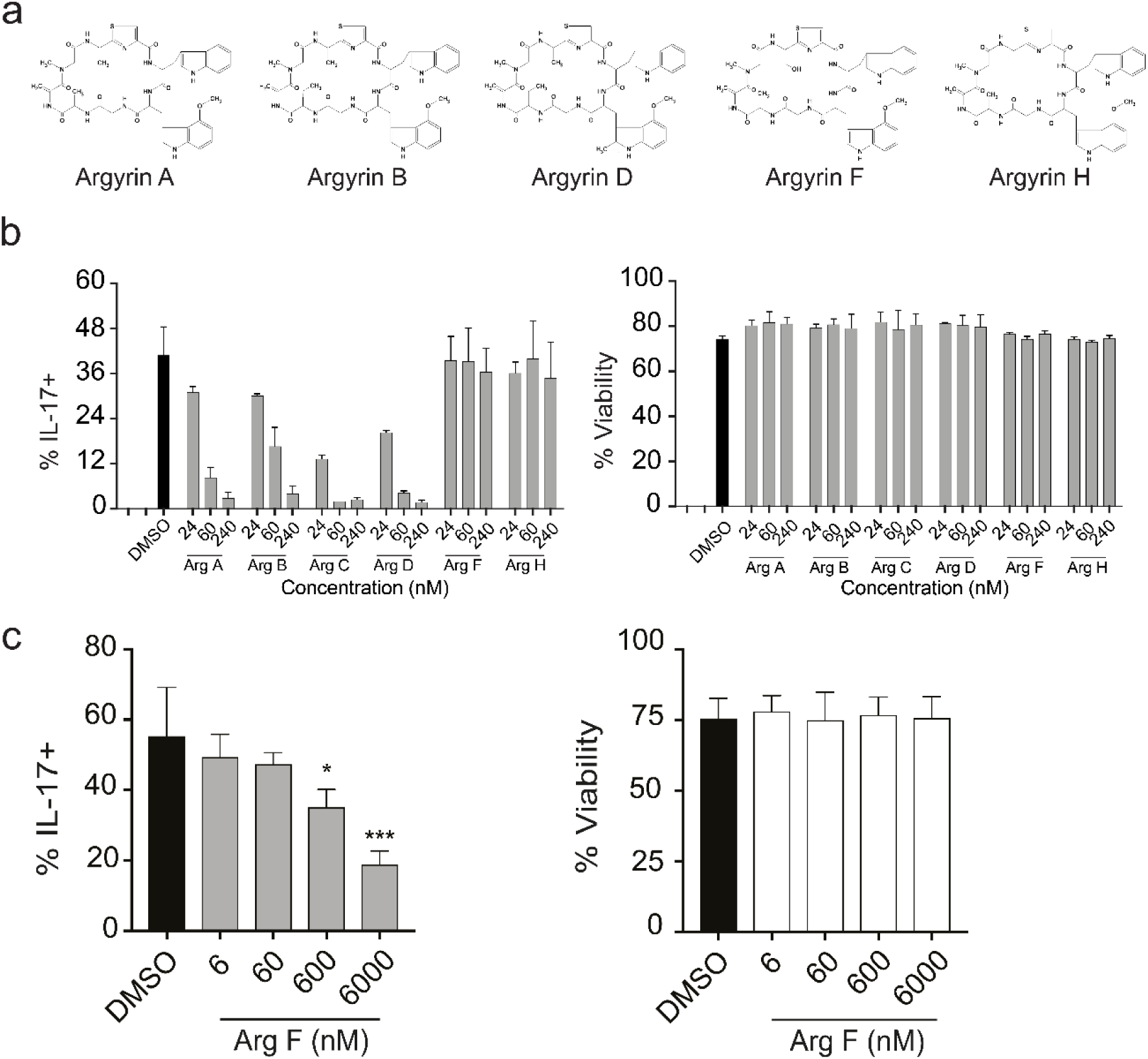
Quantification of the inhibitory effects of Argyrins on Th17 cells, related to Figure 3. **(a)** Chemical structure of Arg family members (A, B, D, F and H). **(b,c)** Mouse naïve CD4^+^ T cells were cultured for 4d under Th17 polarizing conditions in the presence of Arg A, B, C, D, F and H at the indicated concentrations or vehicle (DMSO). Cells were stained for intracellular IL-17A. **(b-c)** Bar graphs show % IL-17A^+^ cells amongst total live CD4^+^ T cells (left) and % viable CD4^+^ cells (right). Results are the pooled means of two **(b)**, three **(c)** individual experiments.

**Figure S2:**
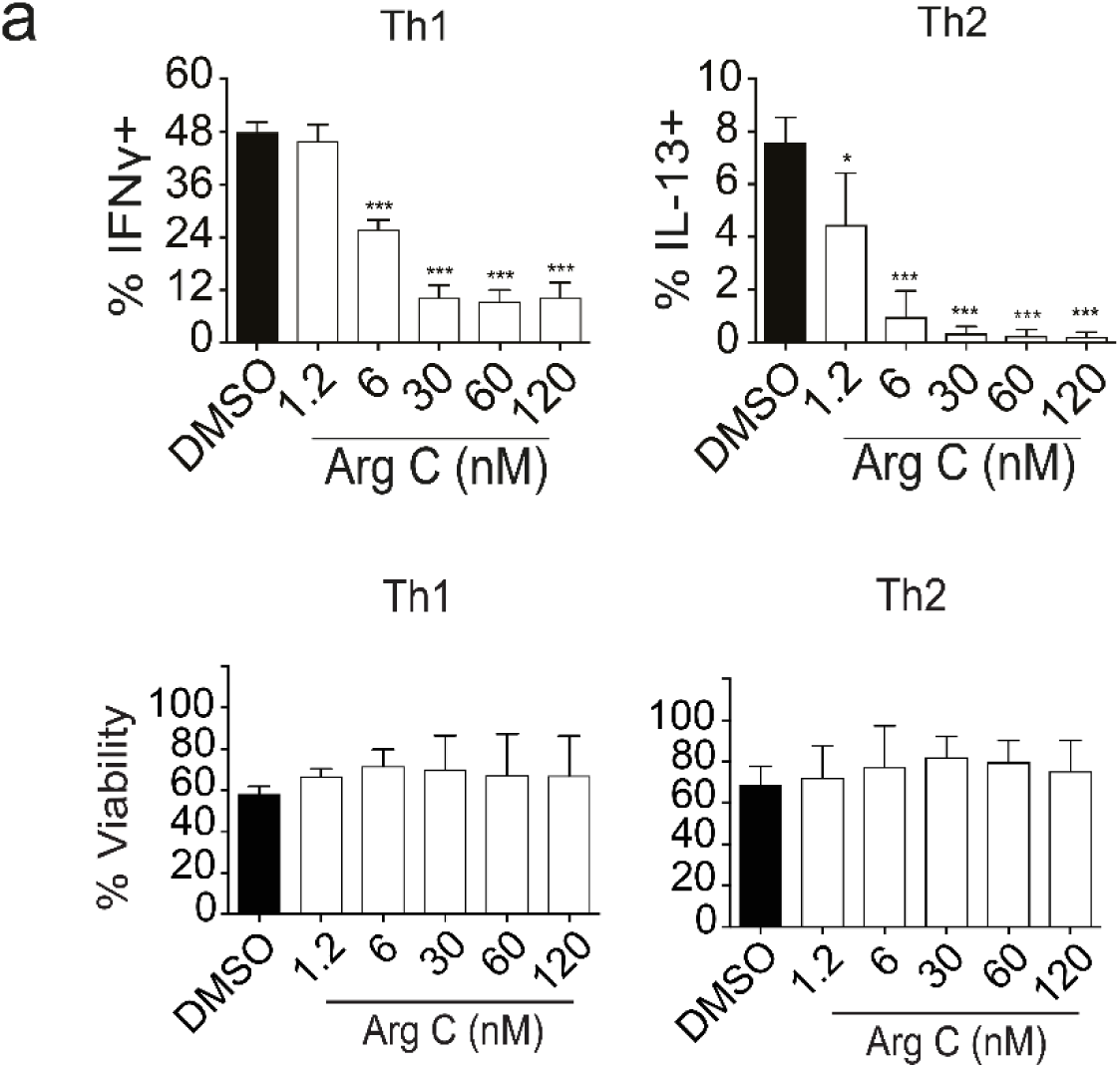
Arg C inhibits Th1 and Th2 effector function, related to Figure 3. Mouse naïve CD4^+^ T cells were cultured for 4d under Th1 and Th2 polarizing conditions in the presence of Arg C (1.2, 6, 30, 60 and 120nM) or vehicle (DMSO). Cells were stained for intracellular IFN-γ and IL-13 and gated on live CD4^+^ T cells. Bar graphs show % of IFN-γ^+^ or IL-13^+^ cells amongst total live CD4^+^ T cells (above) and % viable CD4^+^ cells (below). Results are the pooled means of three independent experiments.

**Figure S3.**
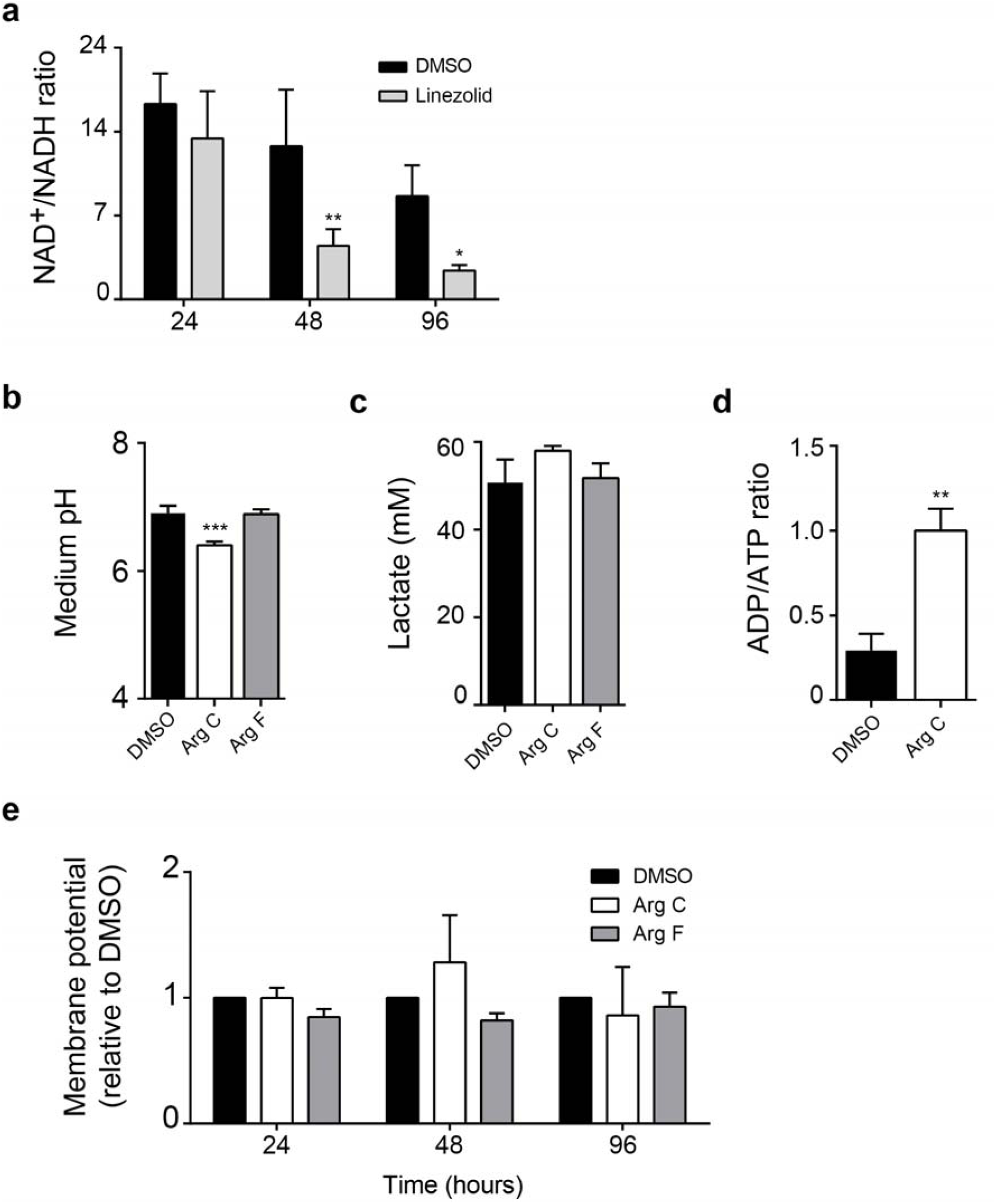
Both linezolid and Arg C treatment impair mitochondria of differentiating Th17 cells, related to Figure 4. **(a-e)** Naïve CD4^+^ T cells were cultured under Th17 polarizing conditions in the presence of indicated drugs Arg C (60nM), linezolid (100µM), Arg F (60nM) or vehicle (DMSO). **(a)** NAD^+^/NADH ratio of CD4^+^ cells treated with linezolid and measured after 24, 48 and 96h of culture. Statistical significance was determined using 2-way ANOVA with Bonferroni multiple corrections test. **(b)** Medium pH, **(c)** lactate concentration and **(d)** ADP/ATP ratio were measured after 96h of culture. **(e)** Mitochondrial membrane potential showing mean fluorescence intensity (MFI) of TMRE incorporation relative to DMSO at 24, 48 and 96h of culture. Results are pooled means of technical replicates from four **(a, b, e)** and three **(c, d)** independent experiments.

**Figure S4.**
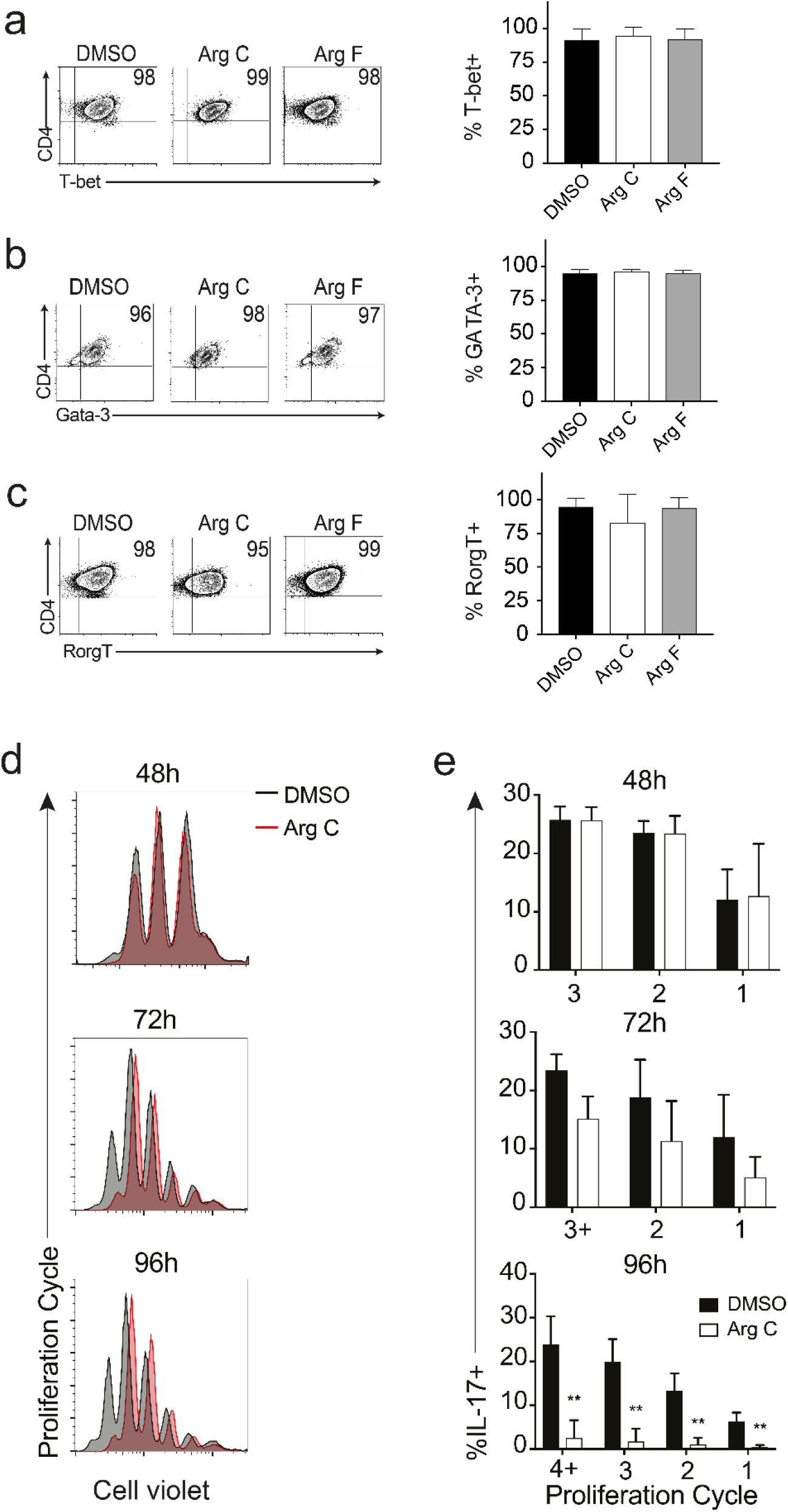
Expression of lineage specific transcription factors is unaffected by Arg C treatment, related to Figure 4. **(a-e)** Mouse naïve CD4^+^ T cells were cultured for 4d under **(a)** Th1, **(b)** Th2, **(c)** Th17 polarizing conditions in the presence of Arg C (60nM), Arg F (60nM) or vehicle (DMSO). Cells were stained for intracellular T-bet, GATA-3 and RORγt and gated on live CD4^+^. **(a-c)** Representative FACS plots showing transcription factor expression (left) and corresponding pooled means (right) **(d,e)** Proliferation was determined using cell proliferation dye CellTrace^TM^ Violet; cells were gated on live CD4^+^ T cells. Representative histograms showing cells gated on live CD4^+^ T cells **(d)**. Cells were stained for intracellular IL-17A and gated on live CD4^+^ T cells. Bar graphs show frequencies of IL-17A^+^ cells in each proliferation cycle at the indicated time point (**e)**. Plots (**a-d**) are representative of three independent experiments and bar graphs (**a-c, e**) are the pooled means of three individual experiments.

**Figure S5.**
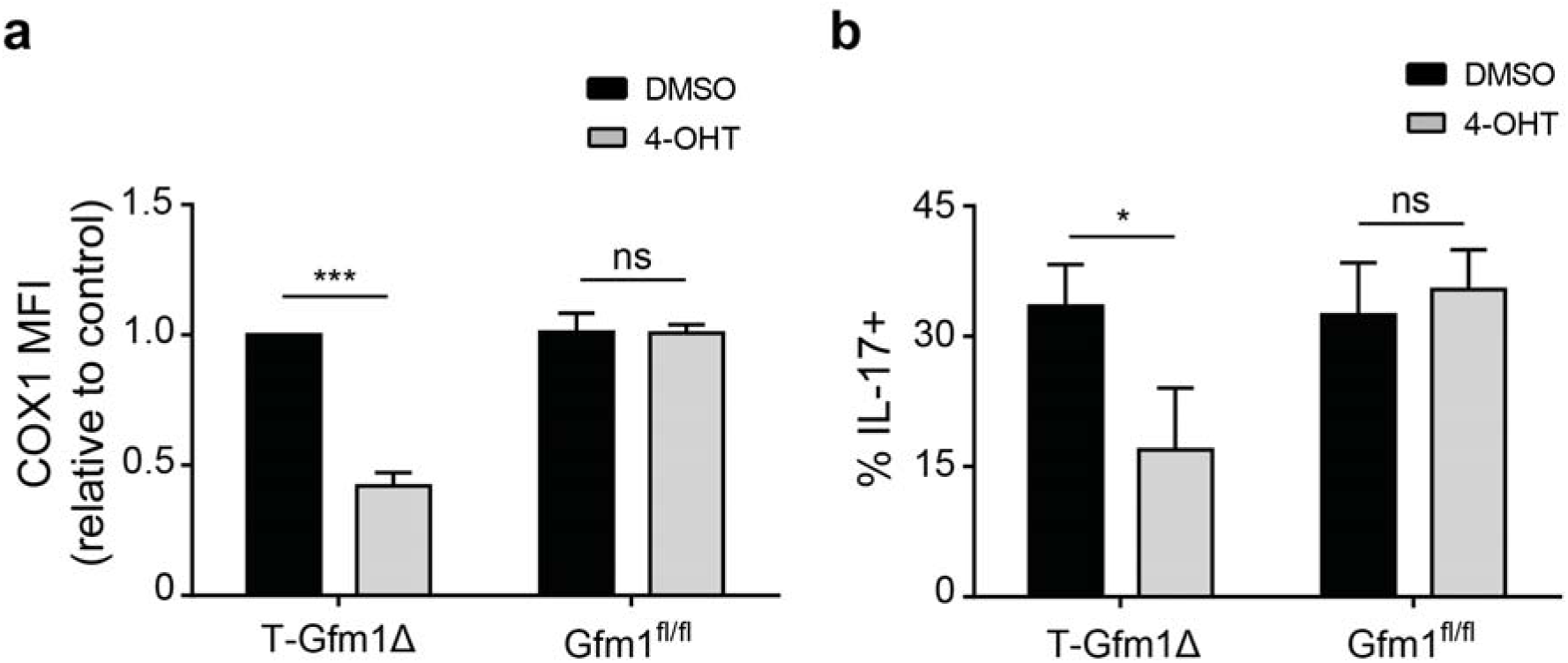
*Gfm1*^flox/flox^ cells differentiate normally in the absence of inducible Cre-activity, Related to Figure 5. Sorted naïve CD4^+^ cells from T-Gfm1Δ mice (Gfm1^flox/flox^Cd4Cre^ERT2/wt^) and littermate controls lacking inducible-Cre activity (Gfm1^flox/flox^Cd4Cre^wt/wt^) were pre-treated with 300nM tamoxifen (4-OHT) or DMSO and maintained in a naïve state for 12 days, prior to being differentiated under Th17 skewing conditions for 4 days. Intracellular levels of COX1 and IL-17A were quantified by FACS. **(a)** Bar graphs shows COX1 pooled and normalized median fluorescence intensity (MFI) relative to DMSO control and **(b)** the pooled means of %IL-17A^+^ CD4^+^ T cells. Bar graphs contain the pooled means from the technical replicates of three **(a,b)** individual experiments.

